## Supplementary Material for "Growth rate-driven modelling suggests that phenotypic adaptation drives drug resistance in BRAFV600E-mutant melanoma"

##### Table of contents

**SM1: Model parameters used in main the article results figures (page 1)**
**SM2: Parameter fitting and complementary simulation results (page 1)**
**SM3: Expressions for effective growth rates under intermittent treatments (page 7)**

### SM1: Model parameters used in the main article results figures

The model parameters used to generate the simulation results shown in the main article are listed in Table S1.

| panel | $\eta_{\text{on}}$ | $\eta_{\text{off}}$ | FM(0,0 nM) | FM(1,0 nM) | FM(0,300 nM) | FM(1,300 nM) | FM(0,500 nM) | FM(1,500 nM) | initial condition |
|---|---|---|---|---|---|---|---|---|---|
| **Figure 2** | | | | | | | | | |
| c; 300 nM; no update | - | - | 0.137 day$^{-1}$ | 0.071 day$^{-1}$ | -0.202 day$^{-1}$ | 0.171 day$^{-1}$ | - | - | Figure 4 (day 1) |
| c; 300 nM; all other panels | 2 | 2 | 0.137 day$^{-1}$ | 0.071 day$^{-1}$ | -0.202 day$^{-1}$ | 0.171 day$^{-1}$ | - | - | Figure 4 (day 1) |
| c; 500 nM; no update | - | - | 0.137 day$^{-1}$ | 0.071 day$^{-1}$ | - | - | −0.202 day$^{-1}$ | 0.171 day $^{-1}$ | uniform |
| c; 500 nM; unbiased | 1 | 3 | 0.137 day$^{-1}$ | 0.071 day$^{-1}$ | - | - | −0.335 day$^{-1}$ | 0.146 day $^{-1}$ | uniform |
| c,e; 500 nM; semi-biased | 3 | 4 | 0.137 day$^{-1}$ | 0.071 day$^{-1}$ | 1 | 3 | −0.268 day$^{-1}$ | 0.171 day $^{-1}$ | left-shifted half normal |
| c,e; 500 nM; biased | - | - | 0.137 day$^{-1}$ | 0.071 day$^{-1}$ | - | - | −0.202 day$^{-1}$ | 0.171 day $^{-1}$ | left-shifted half normal |
| **Figure 3** | | | | | | | | | |
| b; 300 nM | $\eta$ | $\eta$ | 0.137 day$^{-1}$ | 0.071 day$^{-1}$ | -0.202 day$^{-1}$ | 0.071 day$^{-1}$ | - | - | Figure 3a |
| e; 300 nM | $\eta$ | $\eta$ | 0.137 day$^{-1}$ | 0.071 day$^{-1}$ | -0.202 day$^{-1}$ | 0.071 day$^{-1}$ | - | - | Figure 3d |
| b,c; 500 nM | $\eta$ | $\eta$ | 0.137 day$^{-1}$ | 0.071 day$^{-1}$ | - | - | −0.287 day$^{-1}$ | 0.074 day$^{-1}$ | Figure 3a |
| e,f; 500 nM | $\eta$ | $\eta$ | 0.137 day$^{-1}$ | 0.071 day$^{-1}$ | - | - | −0.287 day$^{-1}$ | 0.074 day$^{-1}$ | Figure 3d |
| **Figure 4** | | | | | | | | | |
| no update | - | - | 0.137 day$^{-1}$ | 0.071 day$^{-1}$ | -0.202 day$^{-1}$ | 0.171 day$^{-1}$ | - | - | in top figure row |
| all other panels | 2 | 2 | 0.137 day$^{-1}$ | 0.071 day$^{-1}$ | -0.202 day$^{-1}$ | 0.171 day$^{-1}$ | - | - | in top figure row |

Table S1: Model parameters used to in the simulation results presented in the main article. The initial phenotype distribution (initial condition) is described in the right-most column.

### SM2: Parameter fitting and complementary simulation results

In Figure 2d of the main manuscript, the treatment response models include four parameters that are fitted to match data. These are: $\text{FM}(0, 500\text{ nM}), \text{FM}(1, 500\text{ nM}), \eta_{\text{on}}, \eta_{\text{off}}$. As an alternative approach, one could consider fixing $\text{FM}(0, 500\text{ nM}), \text{FM}(1, 500\text{ nM})$ via log-linear interpolation (the open circles in Figure S1), and thereafter fitting only $\eta_{\text{on}}, \eta_{\text{off}}$. However, with this approach, we were not able to find model parameters that yield simulations which simultaneously match both continuous and intermittent treatment responses. This result is shown in Figure S2, which also demonstrates that for all update models, save the biased one, the 300 nM model and the 500 nM model with four free parameters result in similar and often overlapping simulated treatment dynamics.

In Figures S3 to S6 we have reconstructed Figure 2d (with four free parameters) for different values of $n$, i.e., the number of intermittent phenotypes between the extrema $x = 0$ and $x = 1$, and $\rho$, i.e., the probability that a cell accepts a proposed phenotype update. The results show that the directed (semi-biased and biased) update strategies better capture the cell count dynamics in response to treatments than do the non-directed (no update and unbiased) update strategies for tried values of $n > 0$, given the fitness matrix with linearly interpolated growth rates between the fitted extrema $\text{FM}(0, 500\text{ nM}), \text{FM}(1, 500\text{ nM})$.

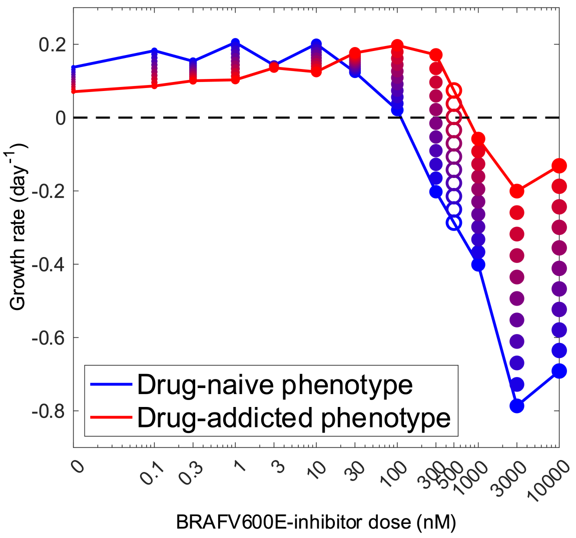


Figure S1: 500 nM growth rates (open circles) can be estimated through log-linear interpolation between 300 nM and 1000 nM growth rates.

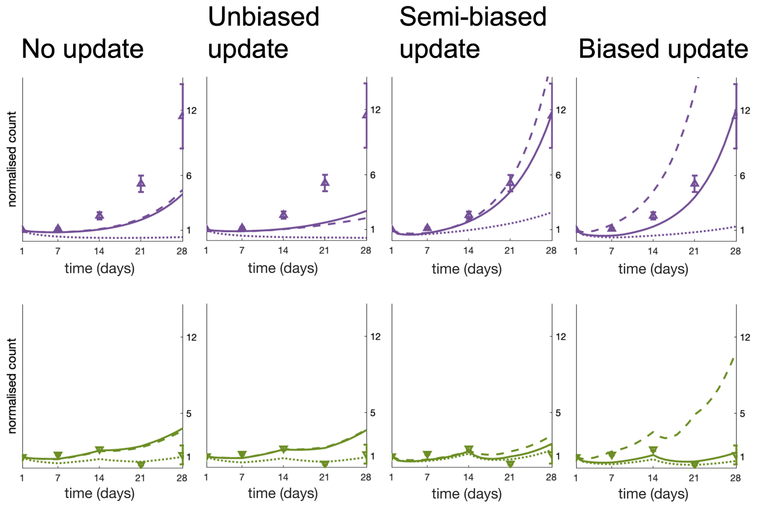


**Continous treatment**

*In vitro* data.

Simulated responses to 300 nM with $\eta_{on} = \eta_{off} = 2$.

Simulated responses to 500 nM with fitted FM(0,500), FM(1,500), $\eta_{on}$, $\eta_{off}$.

Simulated responses to 500 nM with interpolated FM(0,500), FM(1,500) and fitted $\eta_{on}$, $\eta_{off}$.

**Intermittent treatment**

*In vitro* data.

Simulated responses to 300 nM with $\eta_{on} = \eta_{off} = 2$.

Simulated responses to 500 nM with fitted FM(0,500), FM(1,500), $\eta_{on}$, $\eta_{off}$.

Simulated responses to 500 nM with interpolated FM(0,500), FM(1,500) and fitted $\eta_{on}$, $\eta_{off}$.

Figure S2: Simulated responses to continuous (top) and intermittent (bottom) treatments are shown for different phenotype update strategies (left-to-right) and three different model parameter combinations (see legend). In vitro data (triangles) are also shown.

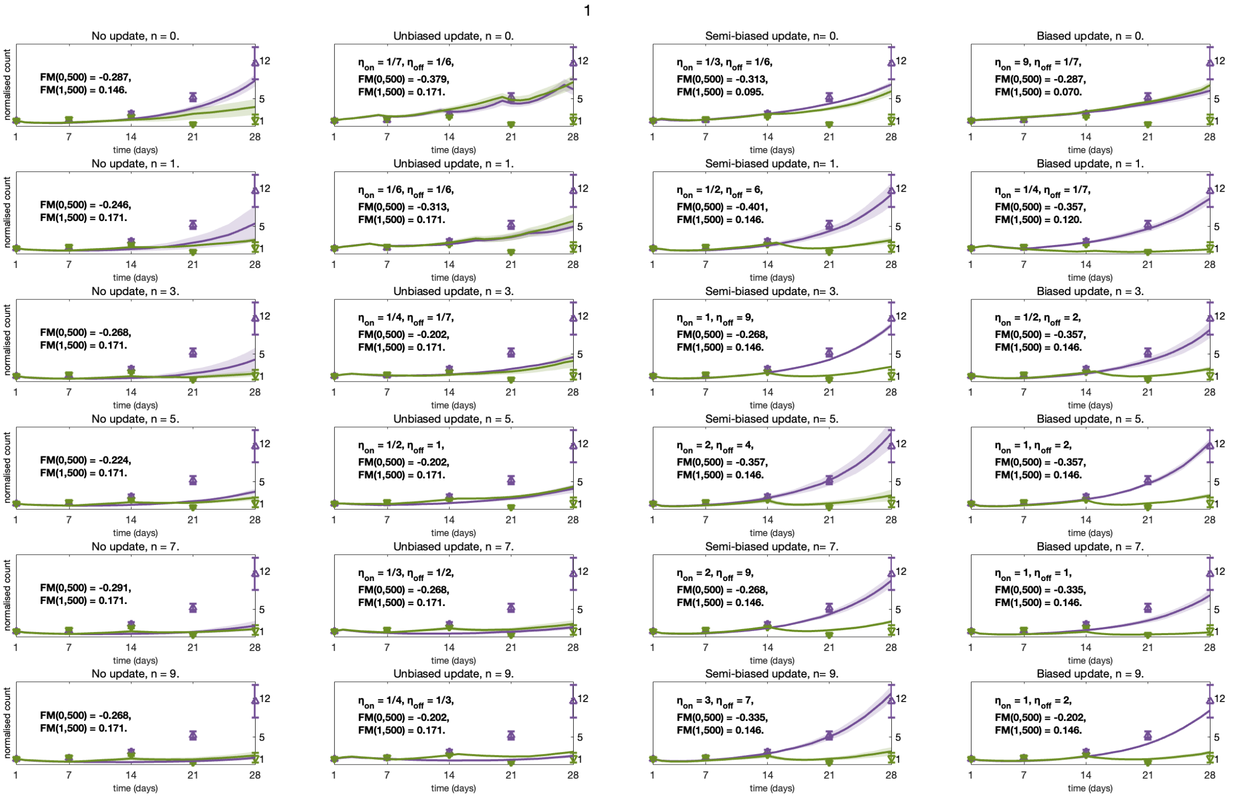


Figure S3: **Simulated treatment responses for $\rho = 1$.** Plots are shown for different phenotype update strategies (left to right), and for varying numbers of intermediate phenotypes $n$ between the extrema $x = 0$ and $x = 1$. The number of phenotypes $n$ increases as we go from higher to lower rows. In each plot, the model parameters $\eta_{\text{on}}$, $\eta_{\text{off}}$, FM($x = 0$,$d = 500$), FM($x = 1$,$d = 500$) that simultaneously minimise the root mean squared error between data and simulations for both continuous (purple) and intermittent (green) treatment responses to 500 nM doses are used.

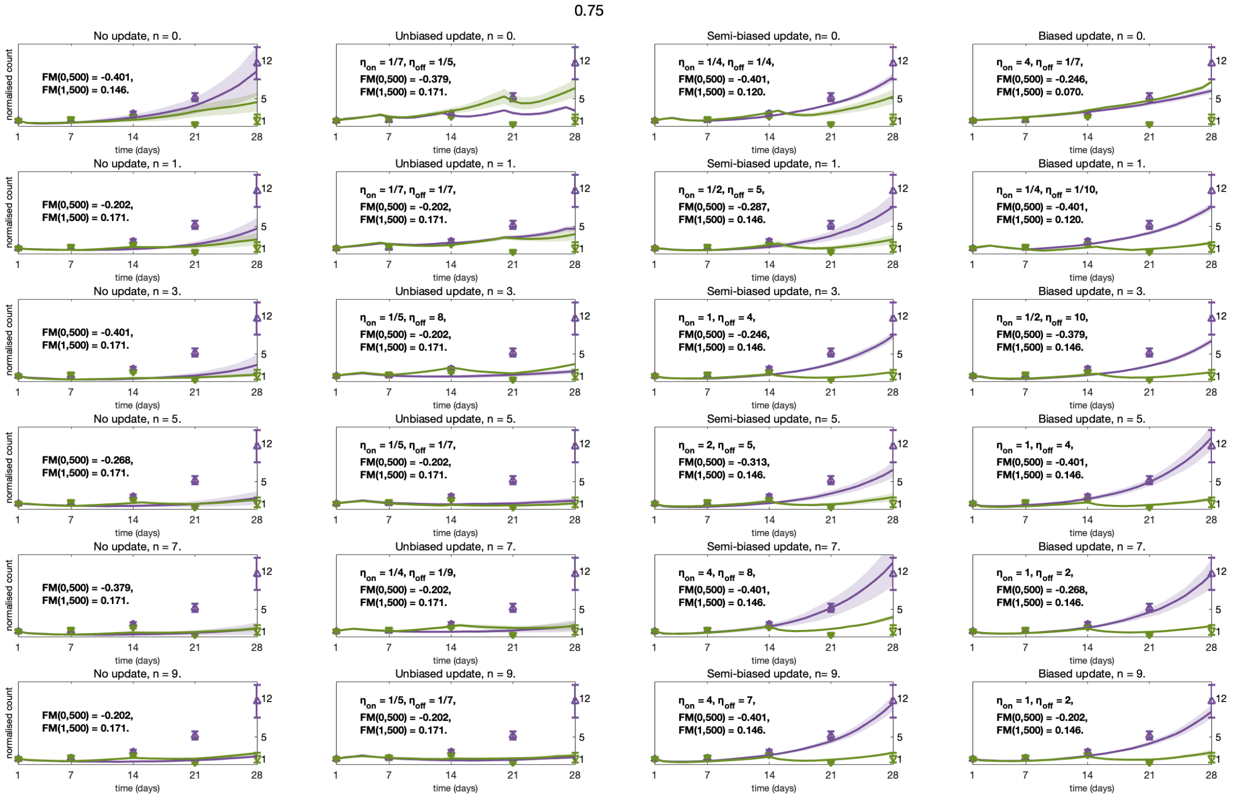

Figure S4: **Simulated treatment responses for $\rho = 0.75$.** Plots are shown for different phenotype update strategies (left to right), and for varying numbers of intermediate phenotypes $n$ between the extrema $x = 0$ and $x = 1$. The number of phenotypes $n$ increases as we go from higher to lower rows. In each plot, the model parameters $\eta_{\text{on}}$, $\eta_{\text{off}}$, FM($x = 0, d = 500$), FM($x = 1, d = 500$) that simultaneously minimise the root mean squared error between data and simulations for both continuous (purple) and intermittent (green) treatment responses to 500 nM doses are used.

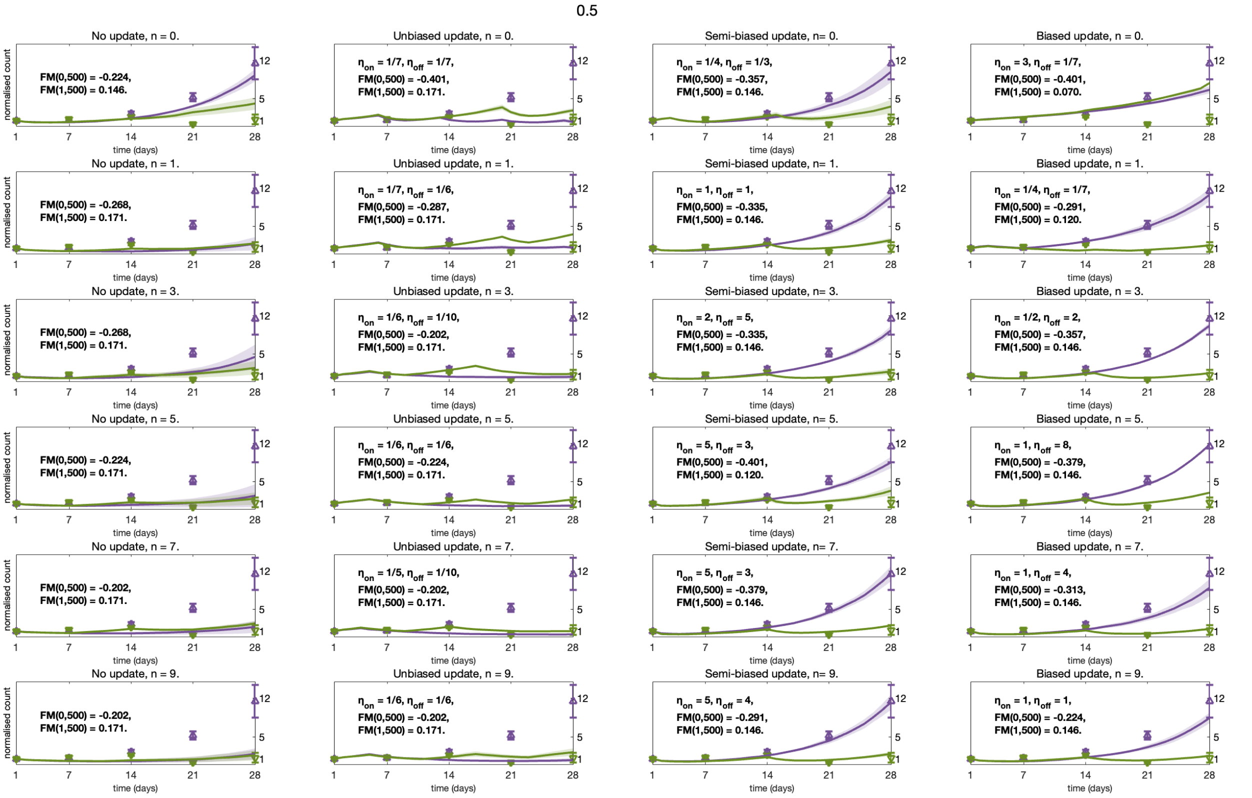


Figure S5: **Simulated treatment responses for $\rho = 0.5$.** Plots are shown for different phenotype update strategies (left to right), and for varying numbers of intermediate phenotypes $n$ between the extrema $x = 0$ and $x = 1$. The number of phenotypes $n$ increases as we go from higher to lower rows. In each plot, the model parameters $\eta_{\text{on}}$, $\eta_{\text{off}}$, FM($x = 0, d = 500$), FM($x = 1, d = 500$) that simultaneously minimise the root mean squared error between data and simulations for both continuous (purple) and intermittent (green) treatment responses to 500 nM doses are used.

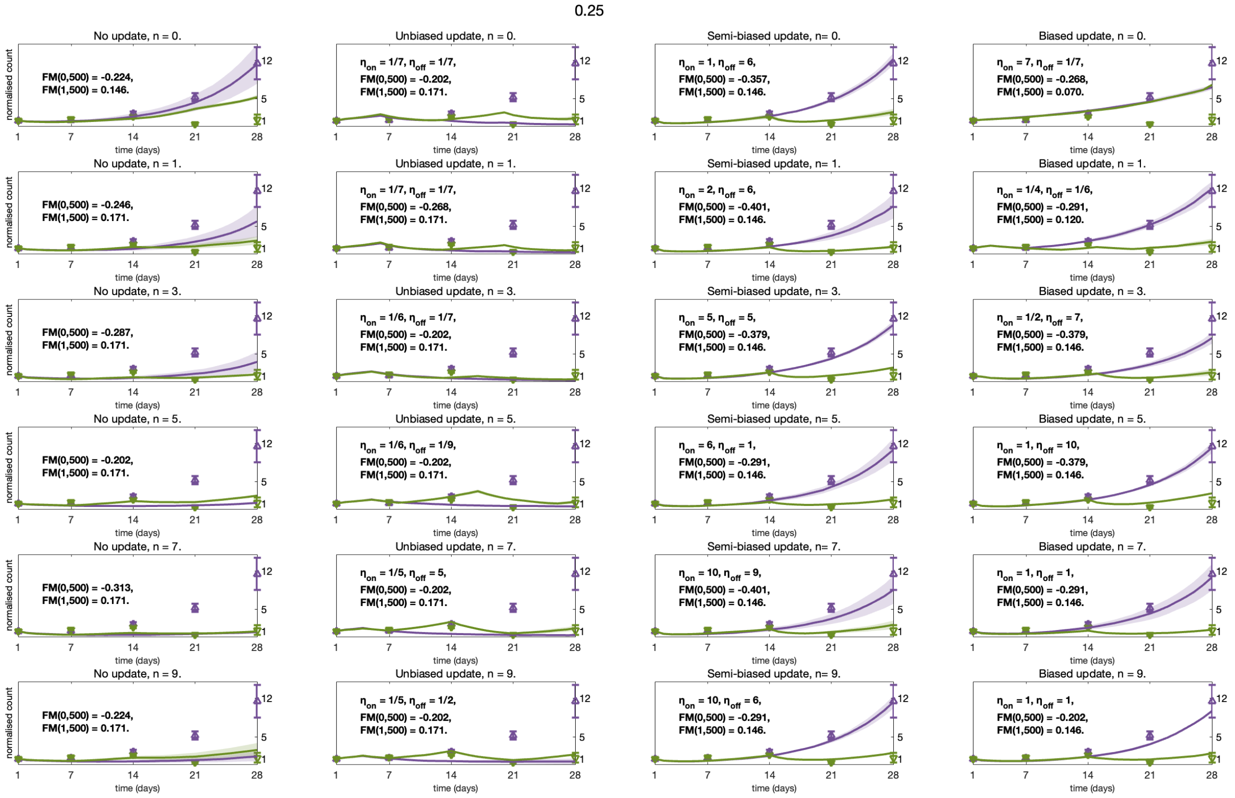


Figure S6: **Simulated treatment responses for $\rho = 0.25$.** Plots are shown for different phenotype update strategies (left to right), and for varying numbers of intermediate phenotypes $n$ between the extrema $x = 0$ and $x = 1$. The number of phenotypes $n$ increases as we go from higher to lower rows. In each plot, the model parameters $\eta_{\text{on}}$, $\eta_{\text{off}}$, FM($x = 0$,$d = 500$), FM($x = 1$,$d = 500$) that simultaneously minimise the root mean squared error between data and simulations for both continuous (purple) and intermittent (green) treatment responses to 500 nM doses are used.

#### SM3: Expressions for effective growth rates under intermittent treatments

A Wolfram Mathematica notebook for calculating long-term effective growth rates for intermittent treatment schedules (Eq. 6 in the main article) are available on the GitHub repository `https://github.com/SJHamis/phenotype_adaptation`. A printed version of the notebook is provided on the next few pages.

### Growth-rate driven modelling study

#### Supplementary material to *Hamis, Browning, Jenner, Villa, Maini, Cassidy (2025).*

In this Notebook we calculate
$\lambda_{eff} = \frac{1}{\tau}\left(\int_0^{t_{on}} FM(x(t), d)\,dt + \int_{t_{on}}^{t_{off}} FM(x(t), 0)\,dt\right)$
for different choices of $t_{on}$ (the duration that the drug is *on*) and $t_{off}$ (the duration that the drug is *off*).

#### Define growth rates as functions of the phenotype state x:

In the notebook, FMxd denotes FM(x,d) where
x=0 is the least drug-adapted phenotype state,
x=1 is the most drug adapted phenotype state,
d=0 means “drug is off”,
d=D means “drug is on” at a selected dose D.

In[1]:=
```
λ0[x_] := FM00 + (FM10 - FM00) x (* no drug *)
λ1[x_] := FM0D + (FM1D - FM0D) x (* drug *)
```

#### State assumptions:

In[3]:=
```
$Assumptions = ω > 0;
```

#### Case 1: $t_{on}, t_{off} > \frac{1}{\omega}$ .

**Drug on:** Phenotypes start at x=0, reach x=1 during climbing period of length $1/\omega$, and then stay at x=1 until time $t_{on}$.
**Drug off:** Cells reach x=0 during a descending period of length $1/\omega$, and then stay at x=0 until time $t_{on}+t_{off}$.
A plot is shown below for (arbitrarily selected) values of $t_{on}$, $t_{off}$, $\omega$ such that Case 1 holds.

In[4]:=
```
x[t_] := Piecewise[{{ω t, t < 1/ω}, {1, 1/ω ≤ t ≤ ton},
    {1 - ω (t - ton), ton < t < ton + 1/ω}, {0, t ≥ ton + 1/ω}}];
Plot[x[t] /. {ω → 0.5, ton → 5, toff → 10}, {t, 0, 15}, AxesLabel → {"t", "x(t)"}]
```

Out[5]=

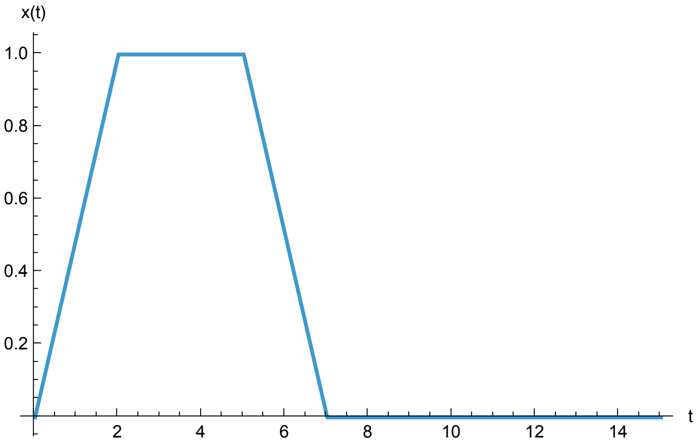


We now symbolically calculate and simplify the expression for the Case 1 integral using $\Delta\text{FM}(d) = \text{FM}(1, d) - \text{FM}(0, d)$.

In[6]:=
```
C1 = 1/τ FullSimplify[
     Integrate[λ1[ω t], {t, 0, 1/ω}] +
     Integrate[λ1[1], {t, 1/ω, ton}] +
     Integrate[λ0[1 - ω (t - ton)], {t, ton, ton + 1/ω}] +
     Integrate[λ0[0], {t, ton + 1/ω, ton + toff}]
   ] /. {FM10 - FM00 → ΔFM0} /. {FM0D - FM1D → -ΔFMD}
```

Out[6]=
$$\frac{\text{FM00 toff} + \text{FM1D ton} + \frac{\Delta\text{FM0} - \Delta\text{FMD}}{2\,\omega}}{\tau}$$

Now customised, numerical values for $t_{\text{on}}$, $t_{\text{off}}$, $\omega$, can be inserted into the expression for C1 to obtain $\lambda_{\text{eff}}$ .

In[7]:=
```
C1num = C1 /. {ΔFM0 → FM10 - FM00, ΔFMD → FM1D - FM0D,  τ → ton + toff};
(*Writing C1 in long format*)
C1num /. {FM0D → -0.2866, FM1D → 0.0741, FM00 → 0.1374, FM10 → 0.07067,
   ω → 0.5, ton  → 5, toff → 10} (*As used in the above plot*)
```

Out[8]= 0.0878047

#### Case 2: $t_{\text{on}} < t_{\text{off}} < \frac{1}{\omega}$ .

**Drug on:** Phenotypes start at x=0, and then reach x=$\omega$t during climbing period of length $t_{\text{on}}$.
**Drug off:** Cells reach x=0 during a descending period of length $t_{\text{on}}$, and then stay at x=0 until time $t_{\text{on}}$+ $t_{\text{off}}$.

A plot is shown below for (arbitrarily selected) values of $t_{on}$, $t_{off}$, $\omega$ such that Case 2 holds.

In[9]:=
```
x[t_] := Piecewise[
  {{ω t, t < ton}, {ω ton - ω (t - ton), ton ≤ t < 2 ton}, {0, 2 ton ≤ t ≤ ton + toff}}]
Plot[x[t] /. {ω → 0.1, ton → 3, toff → 10}, {t, 0, 15}]
```

Out[10]=

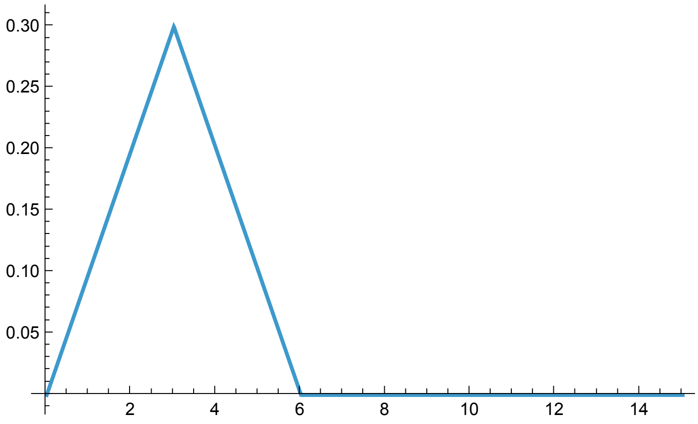


We now symbolically calculate and simplify the expression for the Case 2 integral using $\Delta\mathrm{FM}(d) = \mathrm{FM}(1, d) - \mathrm{FM}(0, d)$.

In[11]:=
```
C2 = 1/τ FullSimplify[
    Integrate[λ1[ω t], {t, 0, ton}] +
    Integrate[λ0[ω ton - ω (t - ton)], {t, ton, 2 ton}] +
    Integrate[λ0[0], {t, 2 ton, ton + toff}]
  ] /. {FM10 - FM00 → ΔFM0} /. {-FM0D + FM1D → ΔFM1}
```

Out[11]=

$$\frac{\mathrm{FM00}\ \mathrm{toff} + \mathrm{FM0D}\ \mathrm{ton} + \frac{1}{2}\ \mathrm{ton}^2\ (\Delta\mathrm{FM0} + \Delta\mathrm{FM1})\ \omega}{\tau}$$

#### Case 3: $t_{off} < t_{on} < \frac{1}{\omega}$ .

**Drug off:** Phenotypes start at x=1, and then reach x=$\omega$t during descending period of length $t_{off}$.
**Drug on:** Phenotypes reach x=1 during climbing period of length $t_{off}$, and then stay at x=1 until time $t_{on}$+ $t_{off}$.
A plot is shown below for (arbitrarily selected) values of $t_{on}$, $t_{off}$, $\omega$ such that Case 2 holds.

In[12]:=
```
x[t_] := Piecewise[{{1 - ω t , t < toff},
    {1 - ω toff + ω (t - toff) , toff ≤ t < 2 toff}, {1, 2 toff ≤ t ≤ ton + toff}}]
Plot[x[t] /. {ω → 0.1, ton → 10, toff → 3}, {t, 0, 15}]
```

Out[13]=

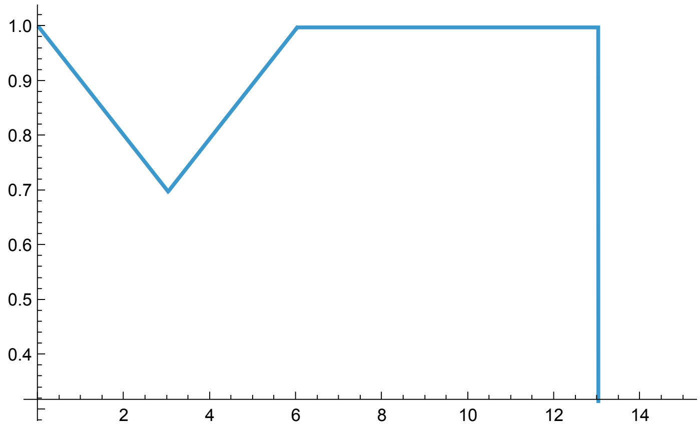

In[14]:=
```
C3 = 1/T FullSimplify[
   Integrate[λ0[1 - ω t] , {t, 0, toff}] +
   Integrate[λ1[1 - ω toff + ω (t - toff)] , {t, toff, 2 toff}] +
   Integrate[λ1[1] , {t, 2 toff, ton + toff}]
  ] /. {FM0D - FM1D → -ΔFMD}  /. {FM00 - FM10 → -ΔFM0}
```

Out[14]=

$$\frac{\text{FM10 toff} + \text{FM1D ton} + \frac{1}{2}\,\text{toff}^2\,(-\Delta\text{FM0} - \Delta\text{FMD})\,\omega}{T}$$

#### Case 4: $t_{on} = t_{off} = \frac{T}{2} = \frac{1}{\omega}$ .

**Drug on:** Phenotypes start at x=0, and then reach x=1 during climbing period of length $\frac{T}{2}$.

**Drug off:** Cells reach x=0 during a descending period of length $\frac{T}{2}$.

A plot is shown below for (arbitrarily selected) values of $t_{on} = t_{off}$ such that Case 2 holds.

In[15]:=
```
x[t_] := Piecewise[{{ω t , t < ton}, {1 - ω (t - ton) , ton ≤ t < 2 ton}}]
Plot[x[t] /. {ω → 1 / 5, ton → 5}, {t, 0, 10}]
```

Out[16]=

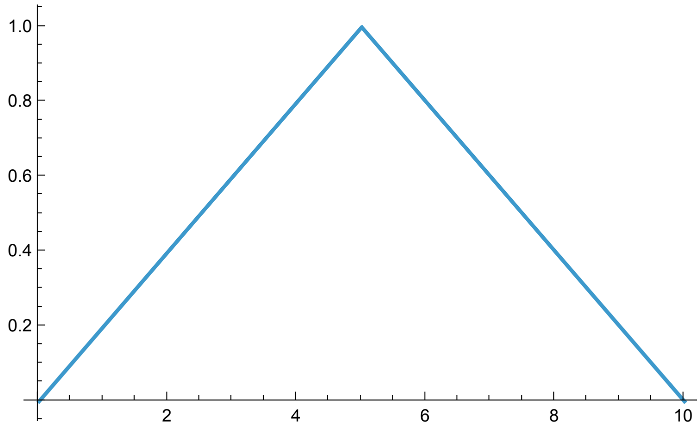

We now symbolically calculate and simplify the expression for the Case 4 integral.

In[17]:=
```
C4 = 1/(2*ton) FullSimplify[
    Integrate[λ1[ω t], {t, 0, ton}] +
    Integrate[λ0[1 - ω (t - (ton))], {t, ton, 2*ton}]
  ] /. {ton*ω → 1} /. {-FM0D - FM10 + 2 (FM0D + FM10) → FM0D + FM10 }
```

Out[17]=

$$\frac{1}{4}\left(\text{FM00} + \text{FM0D} + \text{FM10} + \text{FM1D}\right)$$

We now consider Case 5 and 6 where $t_{on} = t_{off} = \mathrm{T}$.

Case 5: $T < \frac{1}{\omega}$.

Case 6: $T > \frac{1}{\omega}$,

In[18]:=
```
C5 = 1/(2 T) FullSimplify[
    Integrate[λ1[ω t], {t, 0, T}] +
    Integrate[λ0[ω T - ω (t - T)], {t, T, 2 T}]
  ] /. {-FM0D + FM1D → ΔFMD} /. {-FM00 + FM10 → ΔFM0}
```

Out[18]=

$$\frac{1}{4}\left(2\left(\text{FM00} + \text{FM0D}\right) + \mathrm{T}\left(\Delta\text{FM0} + \Delta\text{FMD}\right)\omega\right)$$

In[19]:=
```
C6 = 1/(2 T) FullSimplify[
    Integrate[λ1[ω t], {t, 0, 1/ω}] +
    Integrate[λ1[1], {t, 1/ω, T}] +
    Integrate[λ0[1 - ω (t - T)], {t, T, T + 1/ω}] +
    Integrate[λ0[0], {t, T + 1/ω, 2 T}]
  ] /. {FM0D - FM1D → -ΔFMD} /. {-FM00 + FM10 → ΔFM0}
```

Out[19]=

$$\frac{\left(\text{FM00} + \text{FM1D}\right)\mathrm{T} + \frac{\Delta\text{FM0} - \Delta\text{FMD}}{2\,\omega}}{2\,\mathrm{T}}$$
